# Fast, Efficient, and Precise Gene Editing in the Moss *Physcomitrella patens*

**DOI:** 10.1101/643692

**Authors:** Peishan Yi, Gohta Goshima

## Abstract

Recent years, the bryophyte moss *Physcomitrella patens* has become an emerging model organism for studying conserved signaling pathways and developmental processes during plant evolution. Its short life cycle, ease of cultivation, and high rate of homologous recombination have made it an ideal system for genetic analysis. However, the presence of highly redundant genes and the difficulty of isolating hypomorphic mutants have limited its broader use. Here we developed a simple, fast, and efficient method to generate customized mutants in *P. patens.* We show that transient cotransformation of CRISPR/Cas9 and oligonucleotide templates enables microindel knock-in with high efficiency and accuracy. Using this method, we generated strains carrying various types of mutations, including amino acid substitution, out-of-frame deletion/insertion, splice site alteration, and small tag integration. We also demonstrate that multiplex gene editing can be efficiently achieved to generate putative null and hypomorphic mutants of redundant genes in one step. Thus our method will not only simplify multiple-gene knockout, but also allows the generation of hypomorphic mutants of genes of interest, especially those that are essential for viability.

## INTRODUCTION

The moss *Physcomitrella patens,* one of the basal land plants, has been emerging as a new model organism to address evolutionary changes of plants upon land colonization, such as ancient organ formation and developmental phase transition (Rensing et al., 2008; Prigge and Bezanilla, 2010; Moody, 2019; Sussmilch et al., 2019). Although another basal land plant, the liverwort *Marchantia polymorpha,* has also been developed (Bowman et al., 2017; Ishizaki et al., 2016), several characteristics of *P. patens,* including high regeneration ability, exceptionally high rate of homologous recombination, and ease of long-term time-lapse imaging *in vivo,* have made it a unique system to investigate fundamental questions in plant biology at the molecular and subcellular levels (Schaefer and Zrÿd, 1997; Schaefer, 2001; Cove, 2005; Prigge and Bezanilla, 2010; Vidali and Bezanilla, 2012; Ishikawa et al., 2011; Yamada et al., 2016).

Current functional studies in *P. patens* are mainly carried out by complete gene knockout owing to its high rate of homologous recombination (Schaefer and Zrÿd, 1997; Schaefer, 2001). However, hypomorphic mutants, another valuable source for gene function studies, have been largely ignored. The presence of gene redundancy is the major drawback that limits the characterization of hypomorphic mutants in *P. patens.* Although replacement of redundant genes through homologous recombination to generate knockout and hypomorphic mutants is doable, the entire experiment is laborious and time-consuming, because each gene has to be replaced one by one. RNA interference (RNAi) is a promising tool to overcome this drawback (Bezanilla et al., 2005; Vidali et al., 2007). Targeting highly homologous sequences or using tandem targets allows knockdown of all paralogs in one step (Nakaoka et al., 2012; Vidali et al., 2009). RNAi is also useful for studying the function of essential genes (Vidali et al., 2007). However, construction of stable RNAi lines can be challenging and the knockdown efficiency varies depending on lines (Prigge and Bezanilla, 2010).

The clustered regularly interspaced short palindromic repeat (CRISPR)/Cas9 system is a powerful tool for gene editing, which has been widely applied to many plant species, including *P. patens* (Nomura et al., 2016; Lopez-Obando et al., 2016; Collonnier et al., 2017a, 2017b). By forming a complex, its components, the endonuclease Cas9 and the single guide RNA (sgRNA), are capable of introducing double-strand breaks (DSBs) at predicted genomic loci once the sgRNA is customized with a targeting sequence (Jiang and Doudna, 2017). These DSBs are repaired through the non-homologous end joining or homology directed repair pathways, which leads to subsequent nucleotide changes (Jiang and Doudna, 2017). In *P. patens,* CRISPR/Cas9 has been exploited to generate heritable mutants (Nomura et al., 2016; Lopez-Obando et al., 2016; Collonnier et al., 2017a). However, it is unclear whether precise mutation knock-in can be achieved to generate hypomorphic mutants. In addition, as microhomology-mediated end joining (MMEJ) pathway is active in *P. patens* (Kamisugi et al., 2006; Collonnier et al., 2017a), template-free editing can generate specific deletions at a high frequency, which may not necessarily introduce out-of-frame mutations. For instance, a previous study reported that 74% of the mutations generated by one CRISPR target are identical in-frame deletions (Collonnier et al., 2017a).

Here we report a fast and efficient method to generate customized mutants in *P. patens.* We show that transient cotransformation of CRISPR/Cas9 and oligonucleotide templates induces microindel knock-in at a high efficiency and accuracy. This allows us to introduce various types of mutations into the genome including amino acid substitution, out-of-frame insertion/deletion, splice site alteration, or small tag integration. We also demonstrate that multiplex gene editing can be efficiently achieved, which enables the generation of putative null and hypomorphic mutants of redundant genes in one step.

## RESULTS

To test whether cotransformation of CRISPR/Cas9 and oligonucleotide templates can be used for precise gene editing in *P. patens,* we set up an optimized protocol for vector cloning, transformation, and genotyping (Figure 1). First, the sgRNA plasmid was modified by inserting a drug resistance cassette to facilitate transformant selection. Second, introduction of CRISPR targets was carried out by PCR amplification of the sgRNA vector and subsequent transformation into bacteria without ligation (Figure S1).

**Figure 1.**
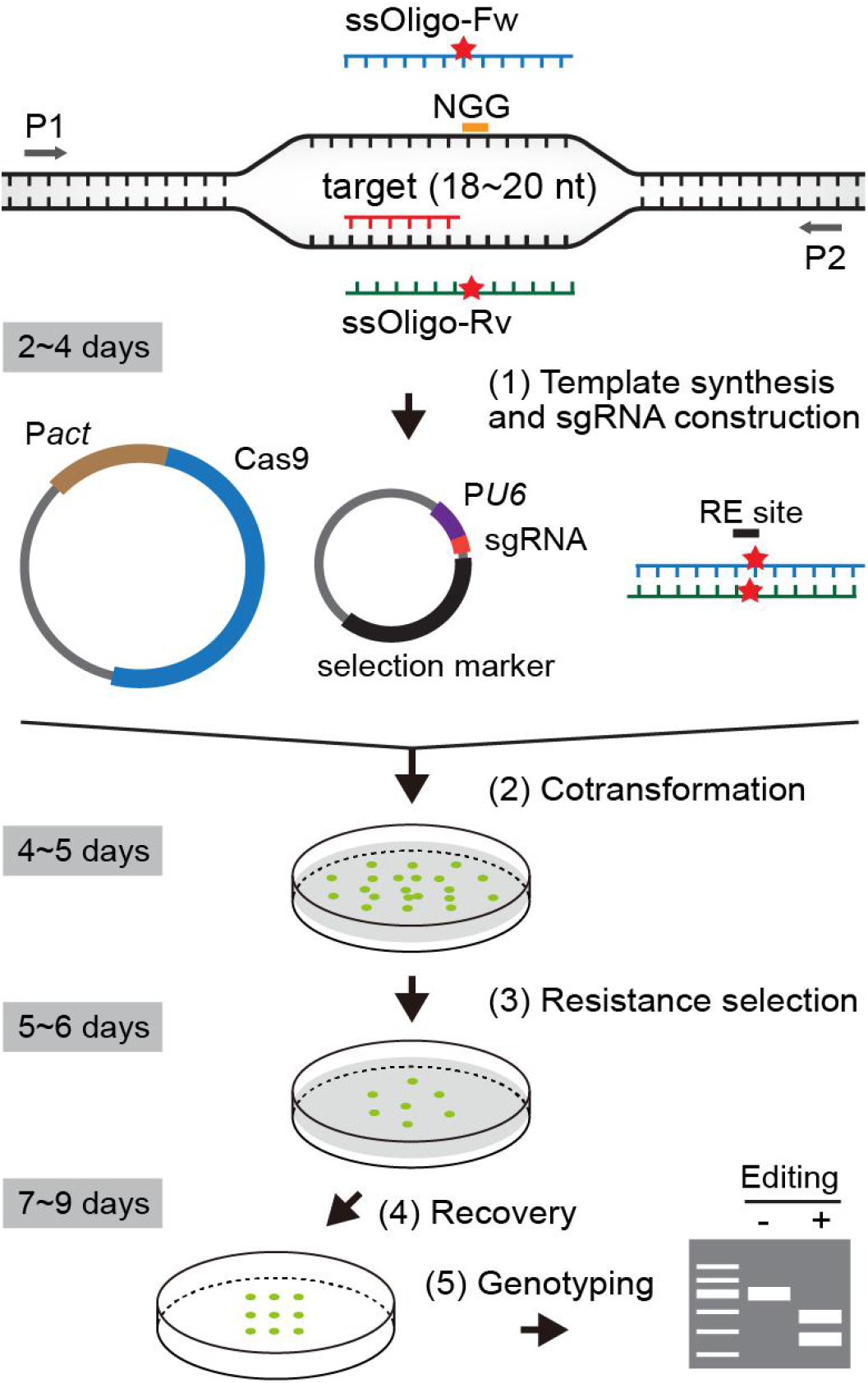
Schematic workflow of precise gene editing in *P. patens.* CRISPR targets are selected at the length of 18-20 nucleotide (nt). An NGG protospacer adjacent motif (PAM) required for Cas9 targeting is shown. Two single stranded complementary oligonucleotide templates (ssOligo) are synthesized at the length of 42-62 nt. The ssOligo in the same direction with CRISPR target is defined as a forward template (ssOligo-Fw) and the complementary ssOligo is a reverse template (ssOligo-Rv). Single stranded templates are annealed before transformation. Red stars, introduced mutations. RE, restriction enzyme site. *PU6, P. patens* U6 promoter. Pact, rice actin 1 promoter. P1 and P2, primers used for genotyping. Approximate time required for each step is indicated on the left.

Third, complementary oligonucleotides at the length of 42-62 nt were synthesized as templates, each containing 20-23 nt microhomology arms (Table S1). Mutations were added between the microhomology arms to introduce a restriction enzyme recognition site and prevent Cas9 cleavage. Therefore, edited lines can be easily identified by restriction fragment length polymorphism genotyping. Fourth, all constructs were transiently transformed into protoplasts, thus enabling fast identification of mutant lines within four weeks.

### Generation of codon substitution mutants

As a proof of principle, we targeted the first exon of *ROP4* (Pp3c10_4950) to mutate its start codon to a stop codon. Two complementary single stranded oligonucleotides at the length of 48 nt were synthesized and designated as forward and reverse templates (ssOligo-Fw and ssOligo-Rv), respectively (Figure 2A). The templates contained the methionine-to-stop (M1stop) mutation and an additional nucleotide change in the nearby 5’UTR to introduce an *XbaI* recognition site (Figure 2A). Double stranded templates (dsOligos) were prepared by annealing the same amount of forward and reverse ssOligos. We transformed the CRISPR/Cas9 plasmids and the templates into protoplasts using the canonical PEG-mediated transformation protocol (Cove et al., 2009; Yamada et al., 2016). After transient resistance selection, we randomly selected four colonies transformed with dsOligos for genotyping and found that two of them were identified by genotyping (Figure 2A & Table 1). Accurate gene modification was confirmed by sequencing. No positive colonies from ssOligo transformation were identified. We then tested another site of *ROP4* to generate a single amino acid substitution W100R with dsOligos (Figure 2B). Genotyping results indicated that 69% of the colonies (N=32) were successfully edited (Table 1). Sequencing of six randomly selected lines showed that five of them were inserted with expected mutations and one contained a 179-bp deletion. Interestingly, one 291-bp deletion mutant was also obtained in the start codon-to-stop substitution experiment (Figure 2A). Analysis of the flanking sequence did not reveal any homology between the joined DNA ends. We speculate that large deletions may occur at the Cas9 cleavage site, therefore preventing homology-directed repair by oligonucleotide templates.

**Figure 2.**
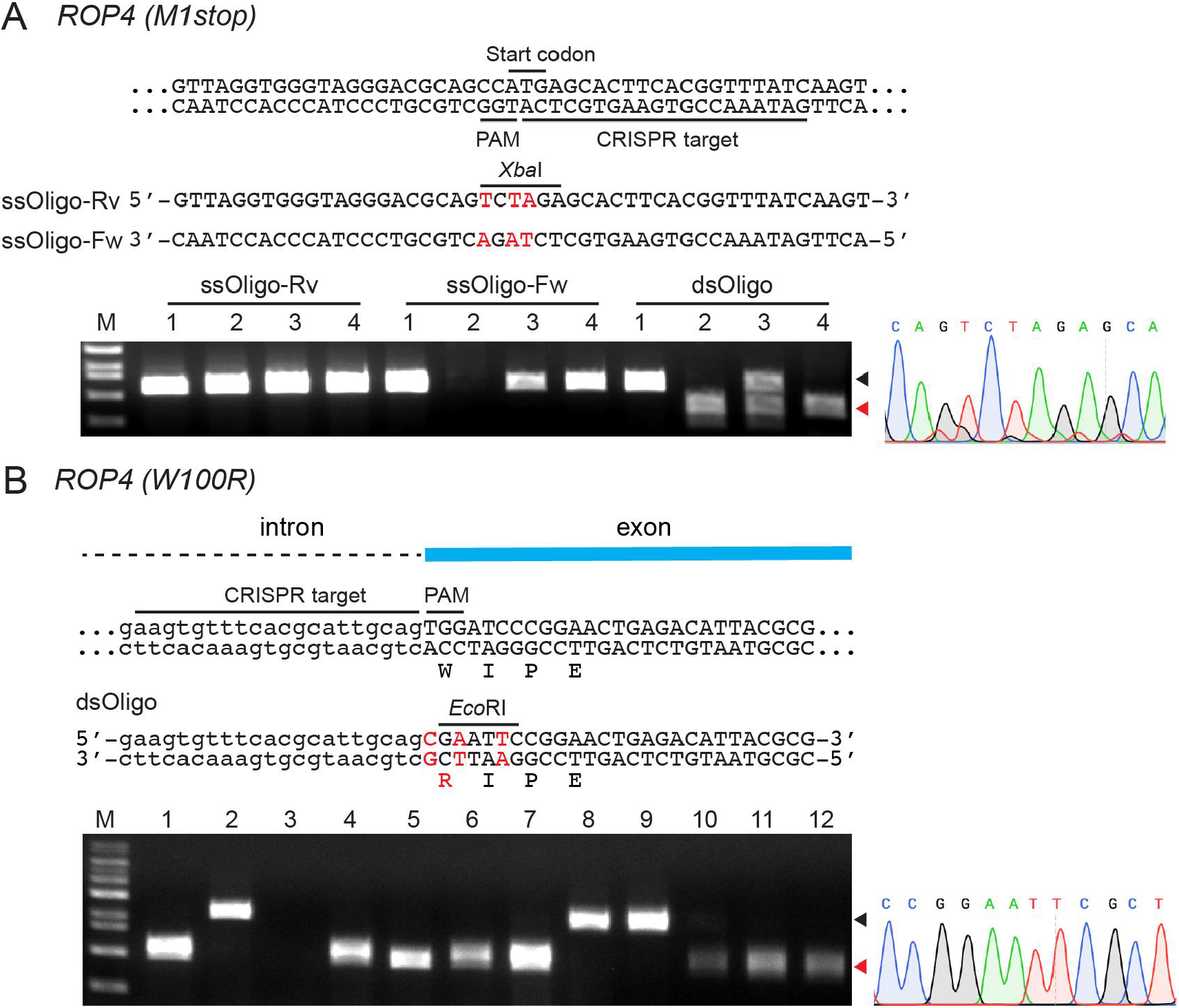
Generation of codon substitution mutants. (A) Substitution of *ROP4* start codon to a stop codon (M1stop). The CRISPR target and PAM motif are underlined. Sequences of forward and reverse single stranded oligonucleotide templates (ssOligo-Fw and ssOligo-Rv) are shown. Mutations are colored in red with an indication of the introduced *XbaI* cut site. The black arrowhead indicates wild-type bands and the red arrowhead indicates putative editing events. In the dsOligo transformation, Line 3 is a chimeric line. Line 2 and 3 are correct knock-in mutants as verified by subculturing and sequencing. Line 4 is a deletion mutant. A representative sequencing result is shown on the right. M, DNA marker. (B) Substitution of the 100th Tryptophan of *ROP4* to Arginine (W100R). Annealed dsOligo template is shown and used for editing. Amino acid change is indicated below the DNA sequences.

**Table 1.**
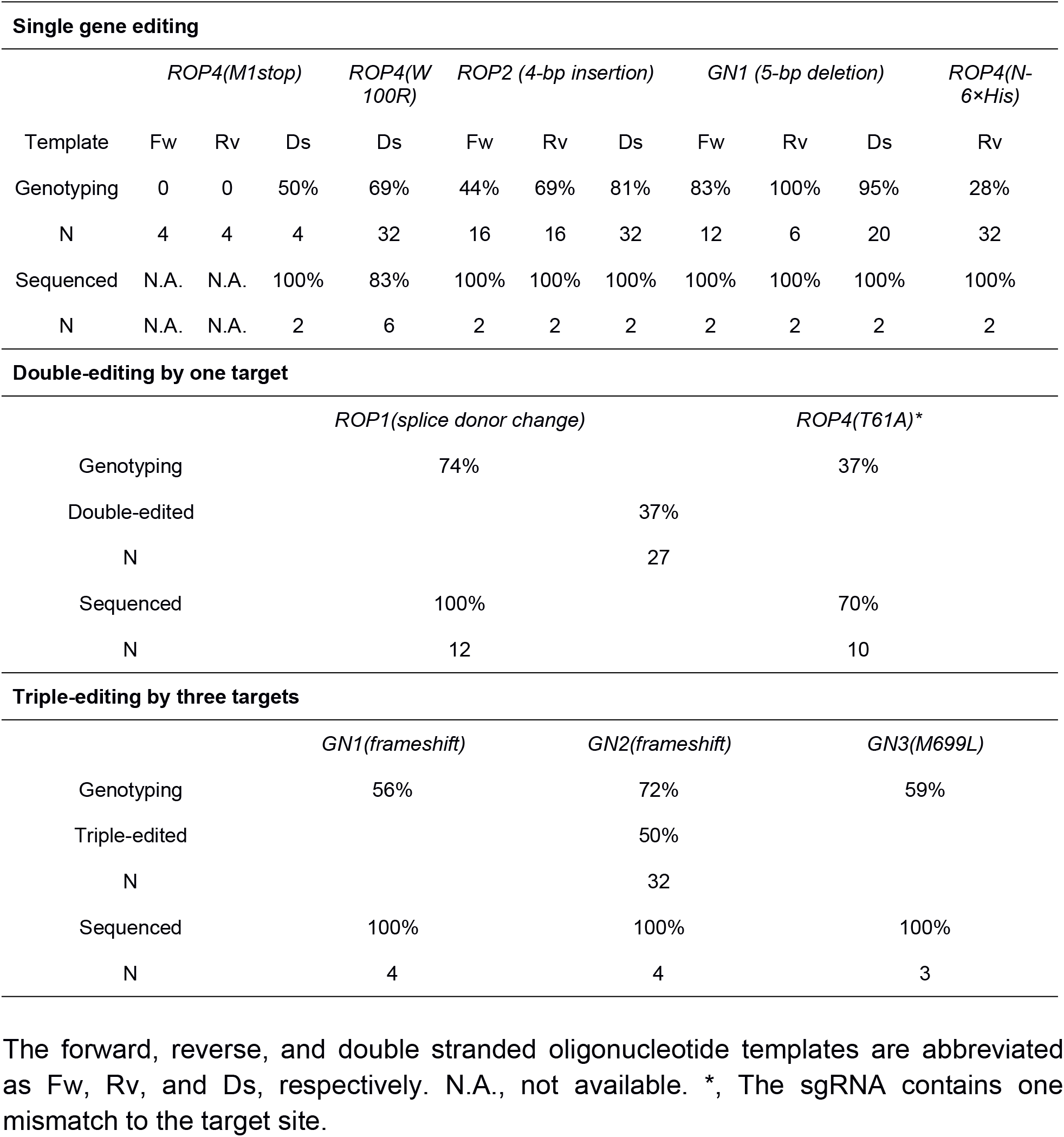
Summary of gene editing results.

### Generation of out-of-frame insertion/deletion mutants

We then tested whether small insertion or deletion can be introduced with Oligo templates. A 4-nt insertion was added to the Oligo templates of *ROP2* (Pp3c2_20700) and a 5-nt deletion was generated for the *P. patens* homologue of *Arabidopsis* Arf-type guanine nucleotide exchange factor *GNOM,* which we named *GN1* (Pp3c9_11830) here (Figure 3A & 2B). Our genotyping results showed that 81% (N=32) and 95% (N=20) of colonies of *ROP2* and *GN1,* respectively, were successfully edited when CRISPR/Cas9 and dsOligos were transformed (Table 1). Interestingly, cotransformation of CRISPR/Cas9 with ssOligo templates of each gene also generated edited mutants (Figure 3A & 3B, Table 1). The editing efficiency of ssOligo template of *GN1* was almost comparable to that of dsOligo template. For *ROP2,* ssOligo template was less efficient. Sequencing of two lines from identified transformants in each category confirmed the correct modification of endogenous loci. To further test the potential of ssOligo in triggering gene editing and the possibility of introducing larger insertions, we transformed a 62-nt ssOligo that targets *ROP4* N-terminus to insert a 6× His tag. 28% of the transformants (N=32) were edited as shown by genotyping (Figure 3C, Table 1) and correct mutations of two lines were confirmed by sequencing. As homologous recombination rate is high in *P. patens* (Schaefer and Zrÿd, 1997; Schaefer, 2001), we repeated the transformation for *ROP2* and *GN1* without supplying Cas9 plasmid to test whether Oligo templates alone are sufficient to trigger gene editing. However, no edited lines were identified when protoplasts were transformed with either ssOligo or dsOligo templates and sgRNA plasmids (N=96 in total). These results indicate that both dsOligo and ssOligo templates can induce mutation knock-in and cotransformation of CRISPR/Cas9 and Oligo templates is necessary for precise gene editing. Our data also suggest that dsOligo templates are generally more efficient than ssOligo templates.

**Figure 3.**
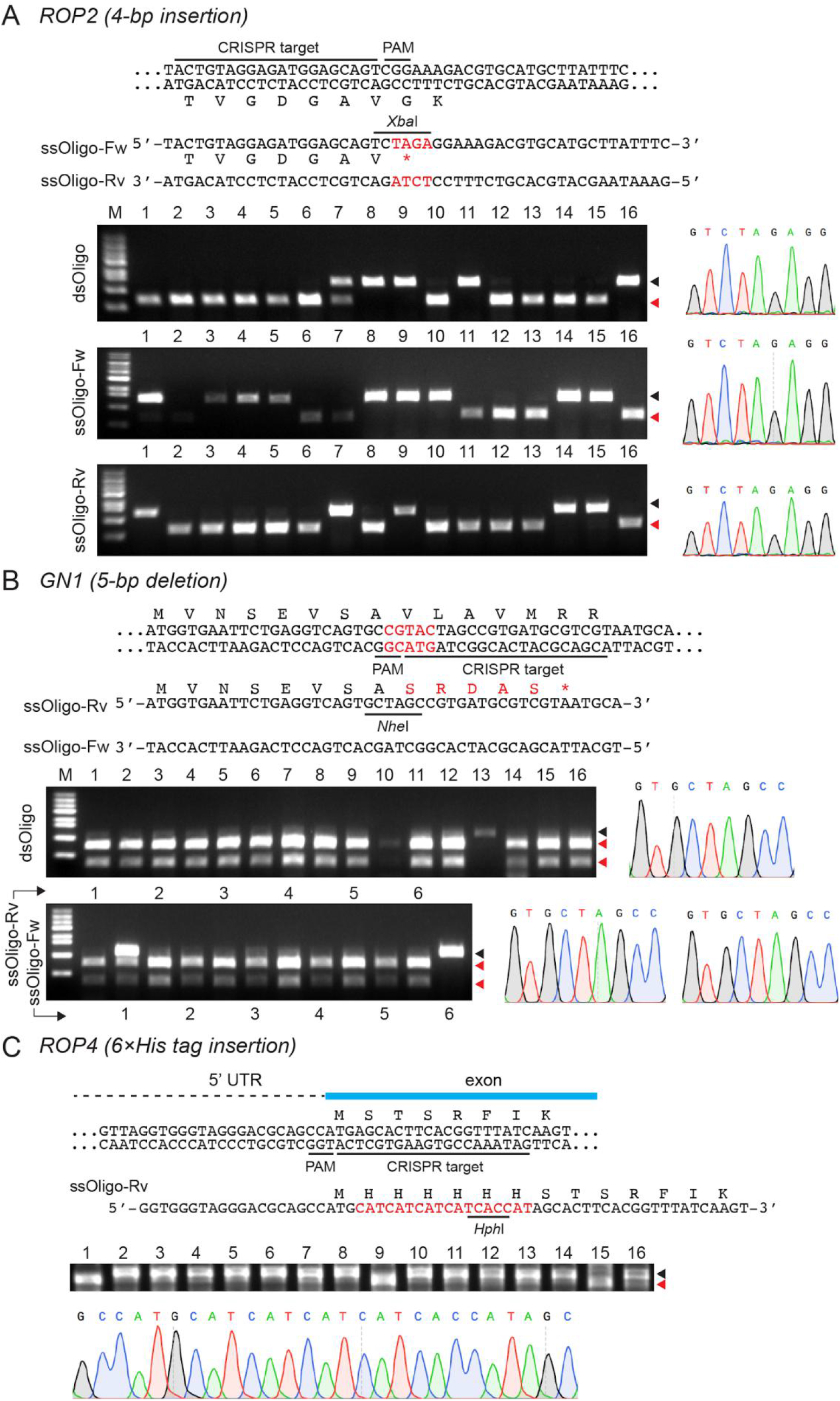
Generation of out-of-frame insertion/deletion mutants. (A) A 4-bp insertion is introduced into the coding region of *ROP2.* Mutations are colored in red with an indication of the introduced XbaI cut site. Amino acid change is shown below the DNA sequences. The genotyping results are shown by a representative gel. The black and red arrowheads indicate the wild-type and edited bands, respectively. Representative sequencing results are shown on the right. M, DNA marker. (B) A 5-bp deletion is generated in the coding region of *GN1.* Mutations of nucleotides and amino acids are colored in red. An *NheI* cut site is introduced after deletion generation. (C) A 18-bp 6×His tag is introduced into the *ROP4* N-terminus. Gene editing was carried out using a 62-nt reverse single stranded template. The His tag insertion is colored in red with an indication of the introduced *HphI* cut site. Translated amino acids are listed above the nucleotide sequences.

### Multiplex gene editing with high efficiency and accuracy

We next investigated the potential of multiple-gene editing and off-target modification. To this end, we selected one sgRNA target of *ROP1,* which also potentially binds to the homologous region of its paralogs *ROP2-ROP4* (Figure 4A). The sgRNA plasmid was cotransformed with the Cas9 vector and dsOligo templates of *ROP1* and *ROP4* that introduce a splice site mutation and a T61A mutation, respectively. Genotyping revealed that 74% (N=27) of the colonies were edited at the *ROP1* locus and 37% were edited at the *ROP4* locus (Figure 4B, Table 1). Interestingly, all of the ROP4-edited mutants were also modified at the *ROP1* site, resulting in a 37% double-editing efficiency (Table 1). This result indicates a strong correlation of editing at both sites, which could be explained by a relatively lower editing efficiency at the *ROP4* locus as a result of one mismatch between the sgRNA and target. To confirm the mutation introduction, we sequenced 12 lines, including the 10 double-edited lines. All of them were correctly edited at the *ROP1* locus and seven of the double-edited lines were correctly edited at the *ROP4* locus. We did not detect any mutations at the *ROP2* or *ROP3* loci. Taken together, our results raise the possibility of targeting multiple sites by a single sgRNA. However, the editing efficiency seems sensitive to target-sgRNA mismatches in *P. patens*. This sensitivity is consistent with previous reports of high specificity of RNA-guided nuclease in template-free editing (Lopez-Obando et al., 2016; Nomura et al., 2016; Collonnier et al., 2017a).

**Figure 4.**
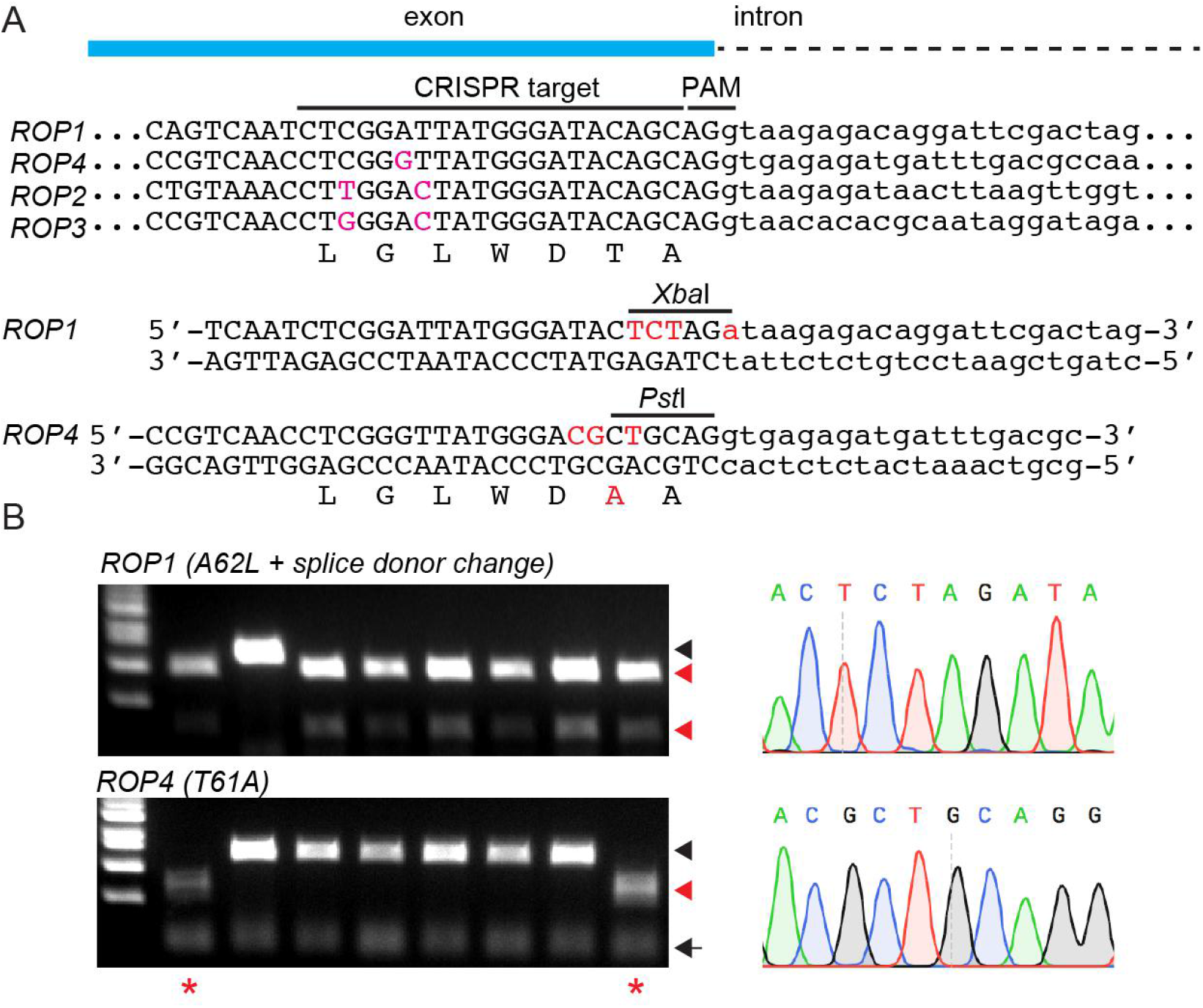
Double-gene editing by one CRISPR target. (A) Alignment of the homologous regions of *ROP1-ROP4.* The CRISPR target and PAM motif of *ROP1* was indicated and used for editing. This target can potentially bind to *ROP2-ROP4.* Mismatch sites in *ROP2-ROP4* are colored in magenta. Two dsOligo templates targeting *ROP1* and *ROP4*, respectively, are cotransformed with the *ROP1* sgRNA plasmid. Mutations in the templates are colored in red with an indication of the introduced XbaI and *PstI* sites. Mutations in *ROP1* result in a Alanine-to-Leucine substitution (A62L) and a nearby splice donor change. Mutations in *ROP4* cause a Threonine-to-Alanine substitution (T61A). (B) Representative DNA gels show the genotyping results at the *ROP1* and *ROP4* loci. Black and red arrowheads indicate the wild-type and edited bands, respectively. Black arrows indicate non-specific bands. Double-edited lines are marked with red stars. Representative sequencing results are shown on the right. M, DNA marker.

We then asked whether cotransforming multiple sgRNAs and Oligo templates would enable multiplex precise gene editing. To test this possibility, we focused on all the three *GNOM* homologues, including *GN1* and the other two paralogs Pp3c15_11320 and Pp3c22_6150, which we named as *GN2* and *GN3,* respectively. CRISPR targets and dsOligo templates were designed to introduce a frameshift mutation in the first exon of *GN2* and a methionine-to-leucine mutation in *GN3* (Figure 5A). The sgRNA cassettes containing *GN2* and *GN3* targets were PCR amplified and inserted into the *GN1* sgRNA plasmid which was used for *GN1* knockout (Figure 3B), resulting in a single vector that expresses sgRNAs targeting all three paralogs. The resultant plasmid was cotransformed with Cas9 vector and three dsOligo templates (Figure 3B and Figure 5A). Our genotyping results showed that 50% (N=32) of the colonies after transient resistance selection were successfully edited at all loci and 28% were subject to a single-gene or double-gene modification (Figure 5B, Table 1). Sequencing analysis of three triple-edited lines and one double-edited line confirmed the correct knock-in of all expected mutations. These results demonstrate that precise editing of multiple genes can be achieved at a high efficiency and accuracy.

**Figure 5.**
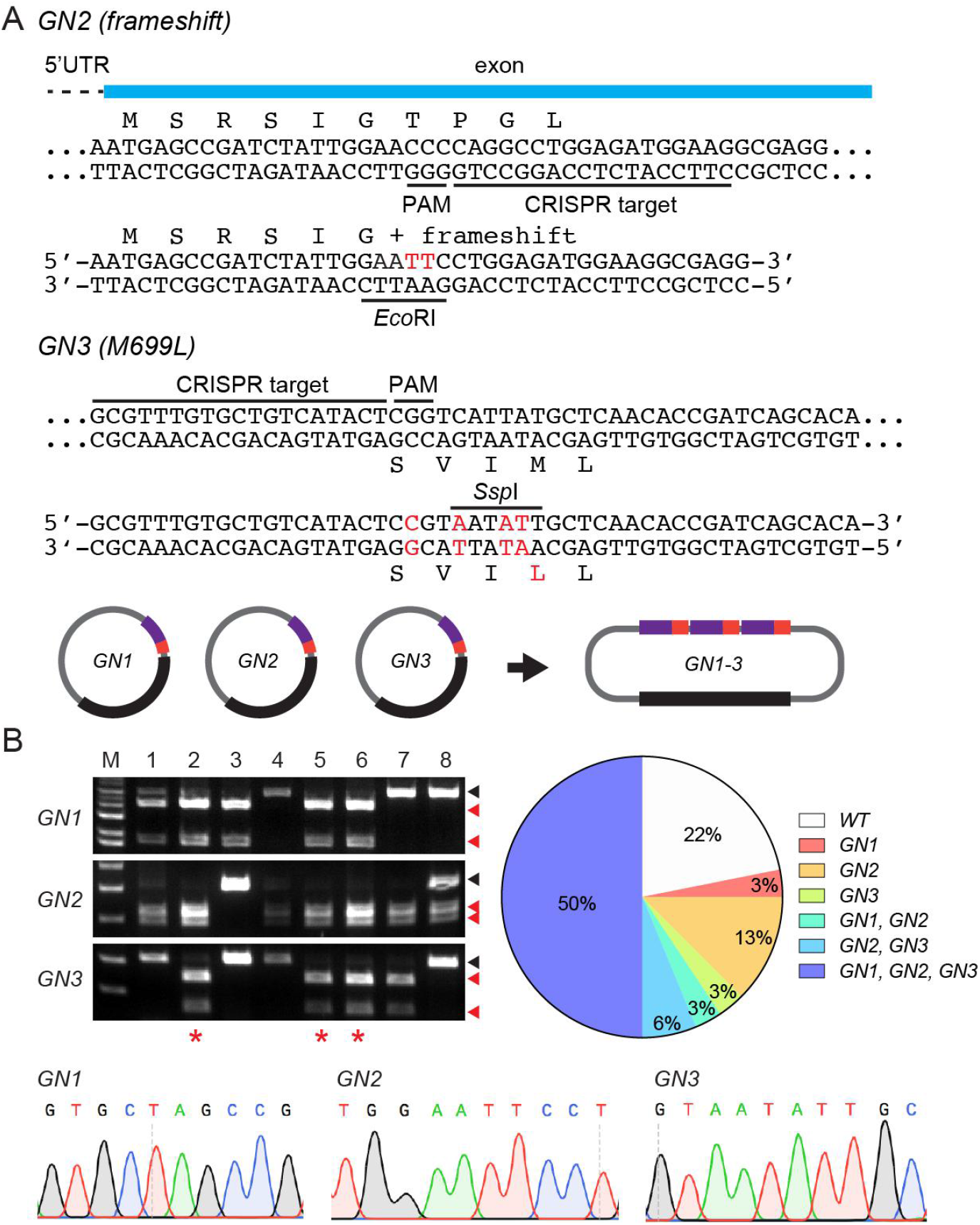
Multiplex gene editing with high efficiency and accuracy. (A) Design of CRISPR targets and templates for multiplex editing. In addition to the constructs used for *GN1* knockout in Figure 3B, CRISPR targets and dsOligo templates targeting *GN2* and *GN3,* respectively, are designed to introduce a frameshift and a Methionine-to-Leucine mutation in each gene. Mutations are colored in red with indications of the introduced *EcoRI* and SspI sites. Amino acid changes are shown above or below the corresponding DNA sequences. A single plasmid that expresses all three sgRNAs was cotransformed with the Cas9 vector and *GN1-GN3* dsOligo templates into protoplasts. (B) Representative DNA gels show the genotyping results at the *GN1-GN3* loci. Black and red arrowheads indicate the wild-type and edited bands, respectively. Triple-edited lines are marked with red stars. Percentage of differently edited lines are shown on the right. Representative sequencing results are shown at the bottom. M, DNA marker.

### *ROP4(W100R)* is a strong hypomorphic mutation

The Rho of Plants (ROP) GTPases are conserved molecules that play diverse roles in cell growth and signaling (Feiguelman et al., 2018). In *P. patens,* four ROP family members have been identified; however, no strong hypomorphic mutants are available (Eklund et al., 2010; Burkart et al., 2015). We constructed the *ROP4(W100R)* mutant on the basis that the mutated residue is highly conserved and its corresponding mutant in yeast is temperature-sensitive (Figure 2B & Figure 6A) (Miller and Johnson, 1997). *ROP4(W100R)* single mutant did not exhibit any obvious phenotypes likely because of gene redundancy. Therefore, we introduced this mutation into the *ΔROP1-3* triple mutant through precise gene editing. As shown in Figure 6B and 6C, *ΔROP1-3; ROP4(W100R)* mutant showed severe growth defects; in contrast, the *ΔROP1-3* mutant exhibited only a moderate growth phenotype. Protonemal cells of *ΔROP1-3; ROP4(W100R)* mutant were dramatically shortened and appeared in a round shape, which is consistent with the observation of strong RNAi knockdown of all ROP genes (Figure 6D) (Burkart et al., 2015). We obtained eight independent lines. All of them showed similar growth defects, indicating that the detected phenotypes were specific to the W100R mutation. We next examined the temperature-sensitive lethality. Wild-type and *ΔROP1-3* mutant strains exhibited slow and retarded growth at 16°C and 28°C, respectively. Although *ΔROP1-3; ROP4(W100R)* mutant grew slower than wild-type and *ΔROP1-3* mutant strains, it was viable under both conditions (Figure 6E). We concluded that *ROP4(W100R)* is a strong hypomorphic mutation and is not temperature-sensitive lethal in *P. patens.*

**Figure 6.**
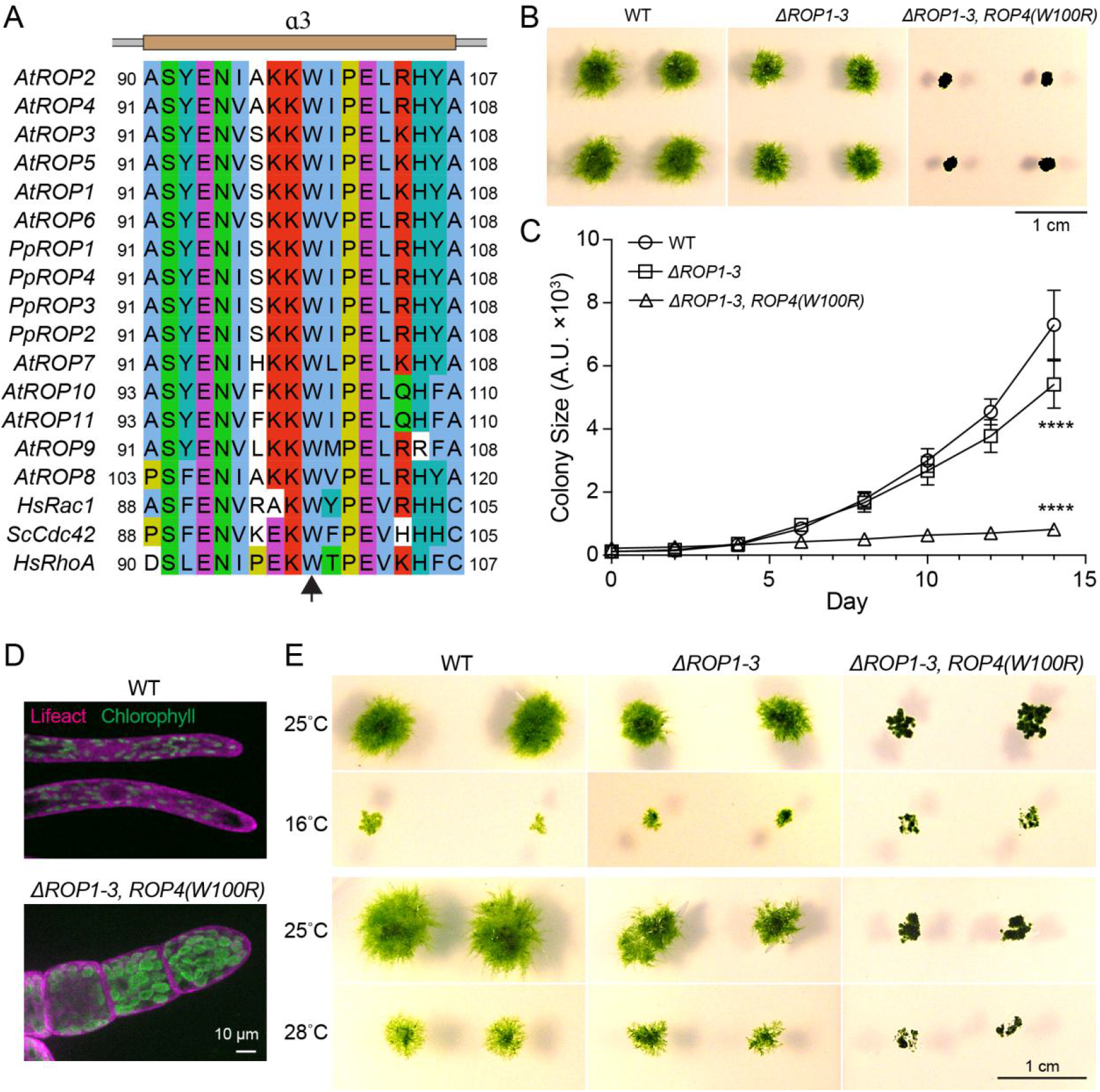
*ROP4(W100R)* is strong hypomorphic mutation and is not temperature-sensitive lethal. (A) Alignment of human RhoA and Rac1, yeast Cdc42, and ROP family members of *P. patens* and *Arabidopsis.* The 100th Tryptophan of ROP4 (arrow) resides in the α3 helix as modeled from the *Arabidopsis* ROP4 structure (PDB:2NTY). (B) Representative images of 14-day old wild-type, *ΔROP1-3,* and *ΔROP1-3, ROP4(W100R)* colonies. Scale bar, 1 cm. (C) Quantification of colony size (mean ± SD, N=16 for each strain). The sizes of 14-day old wild-type and mutant colonies were compared. ****, *p* < 0.0001, two-tailed *t* test. (D) Morphology of protonemal cells of wild-type and *ΔROP1-3, ROP4(W100R)* strains. Scale bar, 10 μm. (E) Representative images of 12-day old wild-type, *ΔROP1-3,* and *ΔROP1-3, ROP4(W100R)* colonies cultured at 25°C, 16°C, and 28°C. Scale bar, 1 cm.

## DISCUSSION

In this study, we present a fast and efficient method to achieve targeted precise gene editing in the moss *P. patens*. This method has several advantages that infer its broad use for genetic studies in the community. First, it allows the generation of various types of mutations including frameshift and hypomorphic alleles, thus can be used for gene knockout and in-depth analysis of any genes of interest, especially those that are essential for viability. Second, compared with canonical gene targeting, it enables multiplex gene knockout within one month. Third, it is a cost-effective and time-wise tool because molecular construction of templates is not required and only transient selection is used for transformation. Moreover, restriction enzyme recognition sites can be easily introduced, which will greatly facilitate genotyping and avoid bulk sequencing. Finally, the ease of obtaining independent stable lines will reasonably ensure the reliability and reproducibility of phenotypic analysis.

Although the CRISPR/Cas9 and oligonucleotide mediated gene editing is highly efficient, several concerns may need to be addressed regarding its use and further improvement. For example, in order to prevent the cut of templates and ensure the length of homologous region, mutations should be close enough to the Cas9 cut site. However, the requirement of an NGG PAM motif limits the modification of specific loci where NGG is not present. A potential solution to this problem is using alternative CRISPR-Cas enzymes or improved Cas9 that recognize different PAM motifs (Adli, 2018; Hu et al., 2018; Nishimasu et al., 2018; Endo et al., 2019; Hua et al., 2019; Ren et al., 2019; Zhong et al., 2019). In our hands, 20-23-bp homology length is sufficient to trigger efficient precise editing. However, increasing homology length might promote editing efficiency further. Because oligonucleotide alone is not sufficient to trigger gene editing, precise editing is very likely triggered by the MMEJ mechanism, and not the canonical homology-directed pathway. MMEJ utilizes 1-16-nt homology sequences to anneal the strands for repair (Sfeir and Symington, 2015). How much the efficiency can be increased by using longer templates is still an open question. Another factor that may affect editing efficiency is the type of mutations and the length of mutated region. Our single-gene editing results suggest a relatively higher editing rate of deletion and insertion knock-in (Table 1). However, longer insert seems disadvantageous as the generation of an 18-bp His tag insertion was less efficient than microindel knock-ins (Table 1). This lower efficiency might also be caused by the use of single stranded templates. Indeed, our data suggest that double stranded templates are more efficient (Table 1), which might benefit from the presence of repair templates of both strands after DSB production.

In general, identified mutants from genotyping exhibit high fidelity of mutation knock-in. However, imperfect mutations were occasionally observed. For example, we obtained deletion mutants from the *ROP4(M1stop)* and *ROP4(W100R)* editing experiments. One mutant from *ROP4(T61A)* editing experiment contained an additional point mutation and two mutants carried complex mutations. It is possible that alternative repair mechanisms are also involved, but occurring at a very low frequency. Another major concern regarding the use CRISPR/Cas9 is potential off-target effects. Previous reports and our data suggest that Cas9 activity seems highly sensitive to sgRNA-target mismatches in *P. patens.* For example, even a single mismatch distant from the seed region yielded a relatively lower editing efficiency (Figure 4 & Table 1) and no off-targets were found when more mismatches existed (Collonnier et al., 2017a; Nomura et al., 2016, 2016). However, in the absence of whole genome sequencing analysis, unexpected off-targeting cannot be simply excluded. Ideally, analyzing independent lines or performing a phenotype rescue experiment would easily solve this problem.

## METHODS

### Moss Culture

Moss strains used in this study were derived from the Gransden strain of *Physcomitrella patens* (Ashton and Cove, 1977). All lines were maintained at 25°C on standard BCDAT plates under continuous light illumination (Yamada et al., 2016) except those that were cultured at 16°C or 28°C for temperature sensitivity analysis.

### Plasmid Construction and Oligonucleotide Synthesis and Preparation

The original Cas9 and sgRNA plasmids are kindly gifts from Fabien Nogué and Mitsuyasu Hasebe (Collonnier et al., 2017a). CRISPR targets were designed using the CRISPOR online tool (http://crispor.tefor.net/) or manually selected at the length of 18–20 nt. For gene knockout, targets were preferentially selected at the first exon. Expression of Cas9 and sgRNA are under the control of rice actin 1 promoter and *P. patens* U6 promoter, respectively. The sgRNA plasmids were modified by adding a selection marker resistant to either Hygromycin B or Nourseothricin. Insertion of CRISPR targets were carried out by using either a ligation method or a site-directed mutagenesis method (Figure S1). In the ligation method, the sgRNA vectors were digested with the type IIS restriction enzyme *Bsa*I (New England Biolabs, cat#R0535S) and dephosphorylated with Calf Intestinal Alkaline Phosphatase (Takara, cat#2250A). CRISPR targets were synthesized in oligonucleotide form and annealed to produce a double stranded fragment which was subsequently ligated to the *Bsa*I treated vectors (Takara, cat#6023). In the site-directed mutagenesis method, a pair of primers carrying parts of the CRISPR targets were used to linearize the sgRNA vectors, yielding DNA products containing 15-bp homology ends (Figure S1). The resultant products were treated with DpnI restriction enzyme (New England Biolabs, cat#R0176S) and transformed into DH5α bacteria without purification. Cas9 and sgRNA plasmids were extracted with the PureLink HiPure Plasmid Midiprep Kit (Invitrogen, cat#K210005). Single stranded salt-free oligonucleotide templates at the length of 42-62 nt were synthesized and dissolved in sterile water at a final concentration of 50 μM. Double stranded oligonucleotide templates were prepared by incubating a mixture of 5 μL each of the forward and reverse oligonucleotides at 95°C for 5 minutes and cooling to room temperature. The sequences of CRISPR targets and oligonucleotide templates were available in Table S1.

### Protoplast Transformation

Circular Cas9 and sgRNA plasmids were cotransformed with oligonucleotide templates into the wild-type strains carrying a Lifeact-mCherry marker or histone H2B-RFP and GFP-tubulin markers (Miki et al., 2016). In general, 10 μg Cas9 plasmid, 10 μg sgRNA plasmid and 10 μL oligonucleotide templates were used appropriately for each transformation. Transformation was carried out using the PEG-mediated protocol as described before (Yamada et al., 2016). In brief, moss tissues were digested with 1% driselase (w/v) in 25-mL 8% mannitol for 30 min. Protoplasts were collected, washed twice with 8% mannitol, and resuspended in the MMM solution (9.1% mannitol, 15 mM MgCl_2_, 0.1% MES pH5.6) at the density of 1.6×10^6^ cells/mL. 300 μL protoplasts were mixed gently with appropriate DNA constructs and an equal volume of PEG solution (40% PEG6000, 10 mM Tris-HCl, 0.1 M Ca(NO3)_2_) and were subsequently incubated at 45°C for 5 min and at 20°C for 10 min. Then protoplasts were sequentially diluted with 300 μL, 500 μL, 700 μL, 1 mL and 4 mL protoplast liquid medium every 3 min and incubated in dark overnight. On the next day, protoplasts were collected, resuspended in 9-mL PRM-T medium, and evenly spread on three cellophane-laid PRMB plates. Cells were cultured for 4-5 days under standard conditions and transferred to BCDAT plates supplemented with 20 μg/mL Hygromycin B or 75 μg/mL Nourseothricin. After 5–6 days, survival colonies were recovered to drug-free BCDAT plates and cultured for another 7-9 days before genotyping.

### Genotyping

A piece of moss protonemal tissue was placed in 30 μL 10× Green PCR buffer (670 mM Tris-HCl pH8.8, 160 mM (NH4)_2_SO_4_, 0.1% Tween 20) and lysed with TissueLyser II (Qiagen, cat#85300). The resultant solution was used as templates for PCR amplification. A genomic DNA fragment containing the CRISPR target site was PCR amplified with primers listed in Table S1 using the KOD FX Neo polymerase kit (Toyobo, cat#KFX-101). Resultant products were digested with 1-2 U restriction enzyme and examined by DNA gel electrophoresis. PCR products used for sequencing were amplified under the same conditions and purified by ethanol precipitation.

### Imaging and Quantification

Moss colonies were imaged with a C-765 Ultra Zoom digital camera (Olympus). Protonemal cells were prepared in glass-bottom dishes and imaged using a Nikon Ti microscope equipped with a CSU-X1 spinning disk confocal scanner unit (Yokogawa), an EMCCD camera (ImagEM, Hamamatsu), and a 40× 1.30 NA lens. A 561-nm and a 488-nm laser lines were used for fluorescence excitation (LDSYS-488/561-50-YHQSP3, Pneum). Image acquisition was controlled by the NIS-Elements software (Nikon). Image processing and quantification of colony size was performed using the Fiji software (https://imagej.net/Fiji). Statistical analysis was performed in Graphpad Prism using twotailed *t* test comparison.

### Sequence Accession and Alignment

Gene sequences are available on the Phytozome website (https://phytozome.jgi.doe.gov). Accession numbers: *ROP1* (Pp3c14_4310), *ROP2* (Pp3c2_20700), *ROP3* (Pp3c1_21550), *ROP4* (Pp3c10_4950), *GN1* (Pp3c9_11830), *GN2* (Pp3c15_11320), *GN3* (Pp3c22_6150). Sequence alignment was performed using the ClustalX software (http://www.clustal.org/).

## Author Contributions

P.Y. conceived the project, performed the experiments, and analyzed the data. P.Y. and G. G. wrote the manuscript.

## Acknowledgments

We thank Fabien Nogué and Mitsuyasu Hasebe for sharing the original CRISPR/Cas9 plasmids. This work is supported by JSPS KAKENHI (17H06471) to G.G. and a long-term postdoctoral fellowship from the Human Frontier Science Program (LT000611/2018-L) to P.Y. The authors declare no conflict of interest.

## Supplemental Figures and Tables

**Figure S1.**
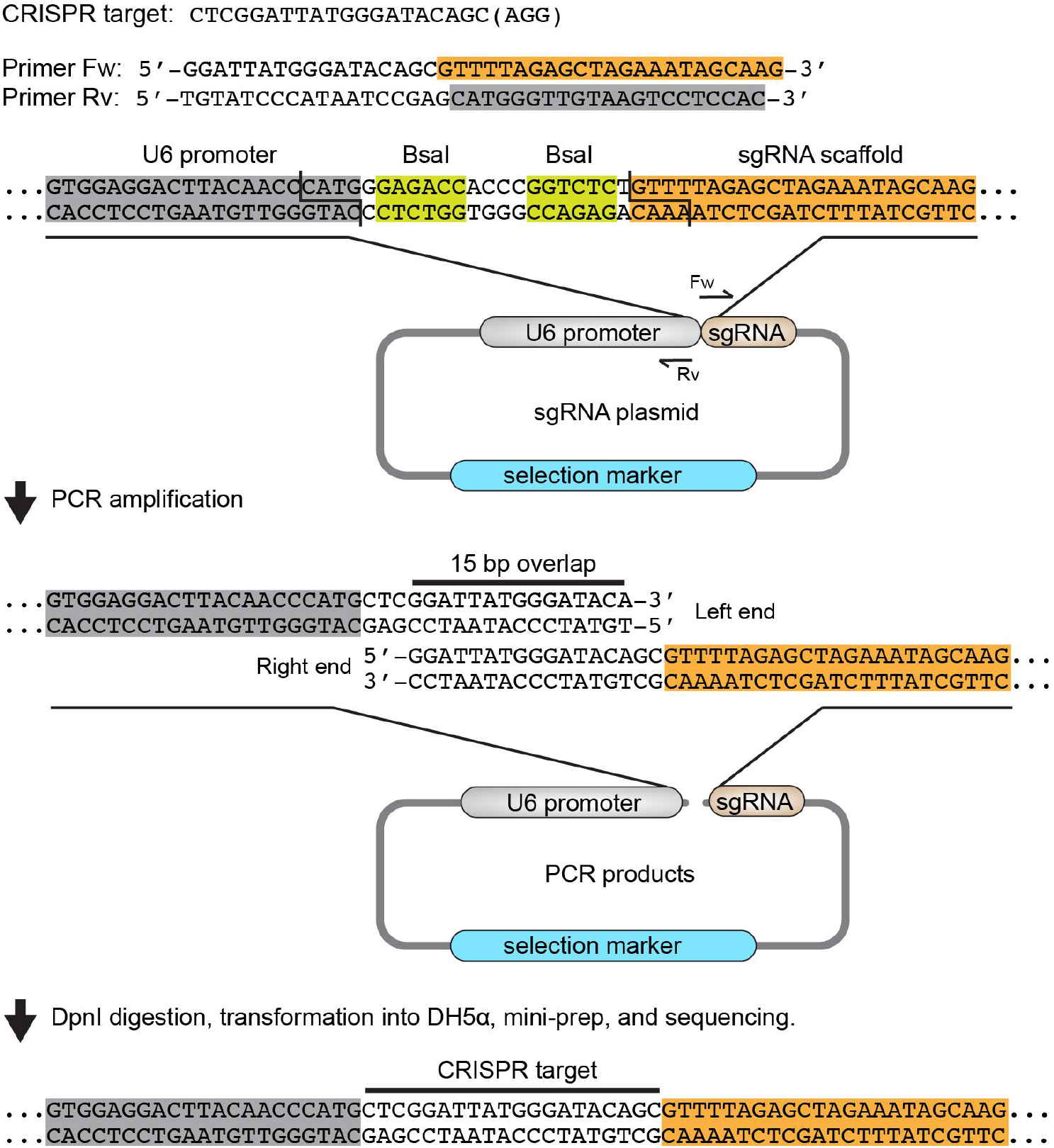
Construction of sgRNA plasmids using the site-directed mutagenesis method. An example of CRISPR targets is presented to illustrate the protocol. A pair of primers, which contain part of the target sequence, were synthesized as Primer-Fw and Primer-Rv to linearize the sgRNA plasmid. PCR amplification yields DNA products containing 15-bp homology sequences at the two ends. The resultant products were treated with *DpnI* and directly transformed into DH5α bacteria without ligation. Spontaneous circularization in the bacteria could efficiently generate correct sgRNA plasmids with inserted targets. This protocol does not rely on ligation treatment and is 100% successful in our hands. As *Bsa*I cut sites (yellow) are reserved between the *U6* promoter and sgRNA scaffold, ligation method can also be used for plasmid construction.

**Table S1. CRISPR targets, oligonucleotide templates, and primers.**

